# Uncertainty and reward histories have distinct effects on decisions after wins and losses

**DOI:** 10.1101/2025.08.14.670176

**Authors:** Shivam Kalhan, Robin Magnard, Yifeng Cheng, Patricia H. Janak

**Affiliations:** Psychological and Brain Sciences, Krieger School of Arts and Sciences, Johns Hopkins University, Baltimore, MD; Solomon H. Snyder Department of Neuroscience, Johns Hopkins University School of Medicine, Baltimore, MD; Kavli Neuroscience Discovery Institute, Johns Hopkins University, Baltimore, MD

## Abstract

Intelligent behavior necessitates an adaptive integration of feedback. It is well-known that animals asymmetrically learn from positive and negative feedback. While asymmetrical learning is a robust behavioral effect, the latent computations behind how animals represent their environments and use this to differentially weight wins and losses is poorly understood. Here we tested whether and how uncertainty and reward histories modulate the weights placed on wins and losses using a behavioral data set collected in rats. We propose a reinforcement learning model that integrates uncertainty history via an unsigned average reward prediction error and a separate subjective reward history component. We showed that in a dynamic probabilistic reversal learning task with blocks of variable reward predictability, ongoing estimation of uncertainty history and reward history both distinctly influenced rats’ sensitivity to wins and losses. In more predictable environments, and under low uncertainty levels, i.e., when rats were certain in making ‘correct’ choices, rats weighted wins more than losses, as indicated by a higher win-stay, and lower lose-shift probability. This asymmetrical learning strategy enabled rats to remain with the correct action, while discounting the influence of rare losses. Further, male rats were more impacted by their reward history, i.e., environmental richness, when making lose-shift decisions, but conversely, female rats were more influenced by their uncertainty history. Hence, we found sex-specific contributions of these latent computations in modulating behavior. We overall demonstrate that asymmetrically weighting wins and losses could form an important behavioral strategy when adapting to ongoing changes in reward and uncertainty history.

## INTRODUCTION

Triumph and defeat are inevitable consequences of an ever-changing environment. Intelligent behaviors necessitate an adaptive integration of these differentially valanced outcomes. It is well-known that animals do not learn symmetrically from wins and losses, but that they generally learn more from positive outcomes, compared to negatives outcomes (Farashahi et al., 2019; Jin et al., 2024; Lefebvre et al., 2017, 2022; Ohta et al., 2021; Palminteri & Lebreton, 2022). Previous work has reasoned that in several contexts, this positivity bias is advantageous to the animal as it makes them less susceptible to unreliable stimuli in their environments which in turn helps them maximize rewards (Cazé & Van Der Meer, 2013; Lefebvre et al., 2022; Palminteri & Lebreton, 2022). However, the environment is constantly changing and requires that the animal keep track of these changes and adapt behavior accordingly. This therefore raises the question of which facets of the changing environmental statistics are relevant to the animal when differentially integrating positive and negative outcomes to adapt behavior.

According to optimal foraging theory (Behrens et al., 2007; Charnov, 1976; Garrett & Daw, 2020; Wittmann et al., 2016), animals track changing reward histories, and use this to make their stay and switch behaviors (Wittmann et al., 2020). Such adaptations allow animals to remain in richer environments. Other work suggests that animals track the changing *uncertainties*, defined as the variability/predictability of the environmental structure (Soltani & Izquierdo, 2019), which could also produce adaptive behaviors by gating learning in proportion to outcome reliability (Le Pelley et al., 2016; Mackintosh, 1975; Pearce & Hall, 1980). However, it is yet to be determined if animals adapt to the changing uncertainty and reward history by asymmetrically weighting wins and losses. It is also currently unclear if uncertainty and reward history play interactive roles or are independently used to adapt these learning strategies. Hence, our overall aim was to investigate whether and how reward history and uncertainty history are dynamically used to influence an animal’s sensitivity to wins and losses. We addressed this aim by using a dynamic probabilistic reversal learning task with blocks of varying reward predictability (i.e., stochasticity). There is also increasing evidence for sex differences in reward guided behaviors in rodents and humans (Aguirre et al., 2024; Chen, Ebitz, et al., 2021; Chen, Knep, et al., 2021; Cox et al., 2023; Lei et al., 2021; Van den Bos et al., 2009), however, it is less understood if distinct influence of uncertainty and reward history states may underlie these behavioral differences. Hence, we tested for the possibility of divergent impact of these environmental statistics on males and females, and whether this may underlie some behavioral differences.

We used computational modelling of behavioral data within the reinforcement learning framework to address these questions (Sutton & Barto, 1998). At the heart of these algorithms is the notion that a surprising outcome generates a learning signal. This learning signal is termed a reward prediction-error (RPE). Importantly, *how much* an animal learns from or weights their RPE is determined by the *learning rate*, which is often a fixed value. Previous work has expanded on the standard reinforcement learning models in an attempt to capture how this learning rate parameter may be dynamically adjusted to capture the differential weighting of RPEs or learning based on the experienced environmental statistics that included aspects of uncertainty and reward rate (Berns et al., 2001; d’Acremont et al., 2013; Grossman et al., 2022; Mackintosh, 1975; Pearce & Hall, 1980; Preuschoff & Bossaerts, 2007; Silvetti et al., 2018; Simoens et al., 2024; Soltani & Izquierdo, 2019; Wittmann et al., 2020). These models greatly increased understanding of uncertainty and reward history gating of learning. However, a critical, yet missing aspect in these models is that the uncertainty history and reward history each may differently influence how much weight is placed on negative and positive feedback, and this additional component could provide mechanistic insights in explaining adaptive behaviors.

To compute an animal’s subjective estimate of their reward history, we used a global reward state (GRS) computation, reflecting the animal’s trial-by-trial estimate of their environmental richness (Grossman et al., 2022; Wittmann et al., 2020). This term captures recency weighted reward history, unlike standard reward averages. Further, to compute an estimate of the animal’s represented uncertainty levels, we computed an unsigned average RPE (avgRPE). The concept of an unsigned avgRPE possibly reflecting an animal’s uncertainty state was introduced by Soltani and Izquierdo (Soltani & Izquierdo, 2019); however, it is yet to be empirically determined how avgRPE may bias an animal’s sensitivity to wins and losses. We hypothesized that the unsigned avgRPE and the GRS computations together may enable subjects to represent relevant facets of their changing environmental statistics, which can then be used to asymmetrically (or symmetrically) gate learning from positive and negative outcomes to adapt behavior.

Our results show that, as expected, subjects do learn more from wins than losses in the probabilistic reversal learning procedure. However, the underlying reasons for this asymmetry vary depending on the distinct subjective reward rates and reward uncertainties present in the environment. Rats used their uncertainty history to down-weight the influence of unreliable losses, and increase the influence of reliable wins, particularly when their environment had a clearer pattern (i.e., less stochasticity). On the other hand, rats were more sensitive to wins when they were in a rich environment (i.e., high reward history state), irrespective of environmental stochasticity (i.e., predictability of rewards) – overall rendering rats less likely to switch from an advantageous action. Further, both males and females showed similar influence of reward history when weighting wins, however, females were less influenced by reward history, compared to males, when making decisions based on losses. Reward history and uncertainty state also interacted to shape decision strategies based on wins, but specifically in males, and in an environment with low stochasticity. These findings together demonstrate that ongoing estimations of uncertainty and reward history differentially impact asymmetrical learning strategies. Because asymmetrical learning strategies change based on highly relevant environmental statistics reflecting likelihood and magnitude of reward, these results suggest that asymmetrical learning is not a maladaptive bias but may form a part of an animal’s adaptive behavioral strategy.

## RESULTS

To address our hypotheses, we analyzed data from our previous study (Cheng et al., 2025; only the non-ethanol exposed rats were analyzed here) where we had trained water-restricted Long-Evans rats (n = 14; 9 males and 5 females) on a dynamic probabilistic reversal learning (dynaPRL) paradigm that had blocks of varying reward probability contrasts between the two actions (Figure 1a-b). There were three different reward probability block types that varied in their reward stochasticity. In the first block type, action one yielded a reward 80% of the time, and action two was rewarded 10% of the time. This was a high contrast block (low stochasticity), as the reward probability difference between the two actions was high. The second block type was a low contrast block (high stochasticity), where action one had a 60% reward probability, and action two was at 30%. Lastly, there was a block with no contrast, with action one and two each having a 45% reward probability (Figure 1b; also see *Methods* for details on the task). These block types were switched within session every 15-30 trials, and male and female rats overall adapted to these reversals by the 6^th^ trial (Figure 1c). Such a design enabled us to investigate how rats used unsigned avgRPE and GRS to differentially weight wins and losses when the environment has a somewhat clear pattern (i.e., in the high contrast block) and when there is a higher level of stochasticity in the environment (i.e., the low contrast and no contrast block).

**Figure 1.**
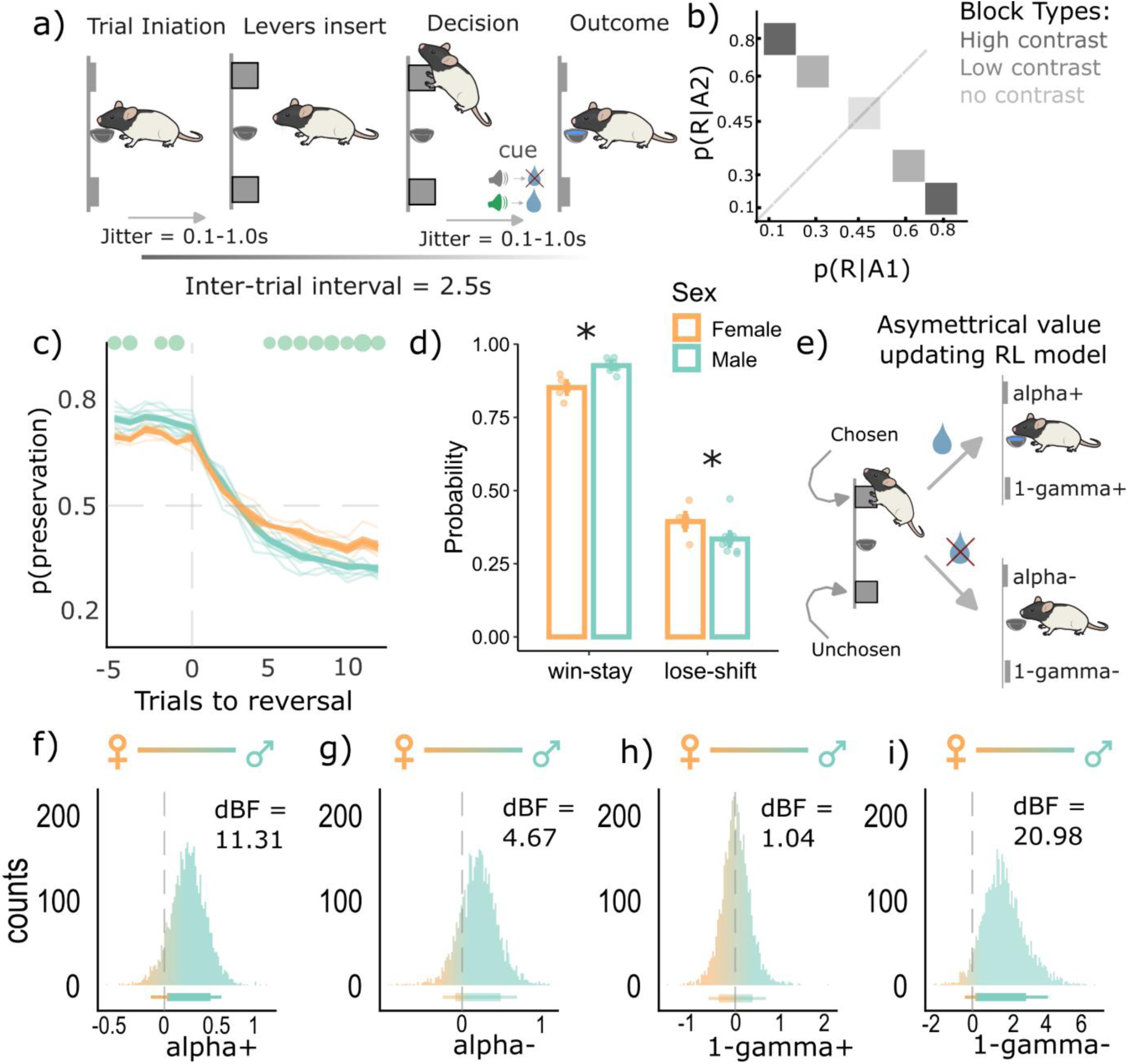
Sex dependent asymmetrical learning behaviors and computations. **a)** Reward choice under changing reward probabilities within the DynaPRL task in which rats initiated trials via magazine entry, following which two levers (left and right) are inserted, and rats made their decision. Outcomes were cued with a clicker noise if rewarded or white noise if unrewarded. The reward was ∼33μL of a 10% sucrose solution. **b)** There were three distinct block types based on their reward probability contrasts. The high contrast block had a p(reward|A1) = 0.8 and p(reward|A2) = 0.1. The low contrast block had a p(reward|A1) = 0.6 and p(reward|A2) = 0.3. The block with no contrast had a 0.45 probability of reward for both actions. **c)** the reversal curve plotted based on p(perseveration) (i.e., likelihood of choosing the previous lever) score for each trial, across all block types. Males and females both reverse by the 6^th^ trial, with females showing a lower probability of repeating the previous best choice (p(perseveration) score) from 5^th^ trial onwards compared to males (green dots are post-hoc tests showing a significant difference between males and females). **d)** Overall win-stay and lose-shift probabilities, across all block types. Both sex groups win-stay more and lose-shift less in our task, however males had a higher p(WS) and a lower p(LS) compared to females. Each dot represents an individual rat, and error bars are standard error of the mean. **e)** Asymmetrical value updating model where chosen and unchosen wins and losses had distinct learning and value decay rates as free parameters **f-i)** Posterior densities from the hyperparameter of sex-group differences for the four value updating parameters from the reinforcement learning model in e. Rightward shifts (above 0) indicate higher parameter values for males compared to females, and the opposite for leftward (below 0) values. Horizontal lines below represent 80-95% highest density interval. Males overall had a higher learning rate from positive (alpha+) and negative (alpha-) outcomes, compared to females. Males also had a higher value decay rate for unchosen action when the chosen action’s outcome was negative (1-gamma-), but similar value decay rates for the unchosen action when the chosen action’s outcome was positive (1-gamma+). *Abbreviations:* dBF = directed Bayes Factor; p(R|A1) = probability of reward given action 1 was taken; p(R|A2) = probability of reward given action 2 was taken. * p < 0.05.

### 1. Rats were more sensitive to wins and less sensitive to losses, and this asymmetrical learning effect was stronger in males compared to females

We first asked whether rats generally had different sensitivities to wins and losses in our task, and whether this effect was sex-specific. To study these win/loss sensitivities, we calculated the overall probability of win-stays (WS) and lose-shifts (LS) for each rat. Rats overall had a higher WS probability compared to the LS probability (Figure 1b, main effect; F(1,24)= 907.28; p = 2.2e-16), indicating that rats were more sensitive to wins compared to losses in our task. Second, there was an interaction between the WS/LS factor and sex (F(1,24) = 14.87, p = 7.6e-04). Subsequent t-tests showed that males had a higher WS probability (p = 0.006, Cohen’s d = 1.70), but a lower LS probability (p = 0.02, Cohen’s d = 1.35), compared to females. This result suggests that males had a greater sensitivity to wins and a lesser sensitivity to losses, compared to females.

To further probe the different win and loss sensitivities in males and females, and the possible computational processes underlying these differences, we used a reinforcement learning model that had four key parameters for updating action value from wins and losses (Figure 1e). The first two were learning rates that updated values for chosen actions from wins and losses (α+ and α−, respectively). The next two were value decay rates for the unchosen actions when the chosen outcome was a win or loss (1-γ^+^ and 1-γ^-^, respectively). Since we were specifically interested in comparing the parameter estimates between males and females, we used group-level hierarchical model fitting (Carpenter et al., 2017; Cheng et al., 2025). Here, the best fit of each rat’s parameters was determined by the group-level (male or female) hyperparameters. The key hyperparameter was the group mean difference, and this gave a posterior density function indicating whether the parameters of interest (learning and value decay rates from wins and losses) were different between the two sex groups, and in which direction (see *Methods*). We found that males had a right-skewed density function for the group mean difference hyperparameter for the learning rate from wins (α+), indicating that males updated more from wins compared to females. The strength of this rightward shift was quantified using directed Bayes Factor (dBF), where males were approximately 11 times more likely (dBF = 11.31) to have a higher α+ than a lower α+, compared to females (Figure 1f). Males were also more likely to have a higher learning rate from negative outcomes (α-; dBF = 4.47; Figure 1g). However, despite having higher learning rates, males had a higher value decay rate from losses (dBF = 20.98; Figure 1i), compared to females, with almost no sex differences in value decay rate for positive outcomes (dBF = 1.04; Figure 1h). These results collectively suggest that a higher value decay from negative outcomes in males may explain their increased sensitivity to wins, but reduced sensitivity to losses, despite having a higher learning rate for both outcomes, consistent with our findings that males have a higher win sensitivity and lower loss sensitivity based on the WS/LS analysis (Figure 1d).

### 2. Asymmetrical learning strategies enabled rats to remain with the ‘better’ high reward probability action by increasing win sensitivity and decreasing loss sensitivity from these actions

Given that rats asymmetrically learnt from wins and losses in our task, and that this effect was sex-specific (Figure 1), we next asked whether rats used asymmetrical learning as part of their strategy to better adapt to the reversals. More specifically, we asked whether rats treated wins and losses differently in the trials immediately post-reversal (early phase; first 6 trials), where they had to learn the new action contingencies, versus the late phase where these action contingencies were most likely learnt (extended from the 7^th^ trial until the end of the block). We found that rats had a higher WS probability in the late phase (p = 0.0001, Cohen’s d = 2.04), compared to early, with no differences in LS probabilities between phases (p = 0.44, Cohen’s d = 0.17), see Figure 2a. Rats were therefore more sensitive to wins in the late phase of a given block. We reasoned that developing a strategy of increasing the WS probability in the late phase may serve the rat to remain with the ‘better’ high reward probability action, while also reducing the chances of incorrectly switching to the ‘worse’ low reward probability action based on rare losses. If this was indeed the case, we should find that the better actions (compared to worse actions) have a higher WS probability and a lower LS probability in the late phase. We therefore calculated WS and LS probabilities separately for ‘better’ and ‘worse’ actions (Figure 2b left) in the late phase. We found that better actions did have a higher WS probability compared to worse actions (main effect of action type: F(1,36) = 57.53, p = 5.68e-09; Figure 2c-d). Further, better actions also had a lower LS probability (F(1,36) = 436.29, p = 2.20e-12; Figure 2e-f). Such a strategy could increase the likelihood of rats persisting with the better option.

**Figure 2.**
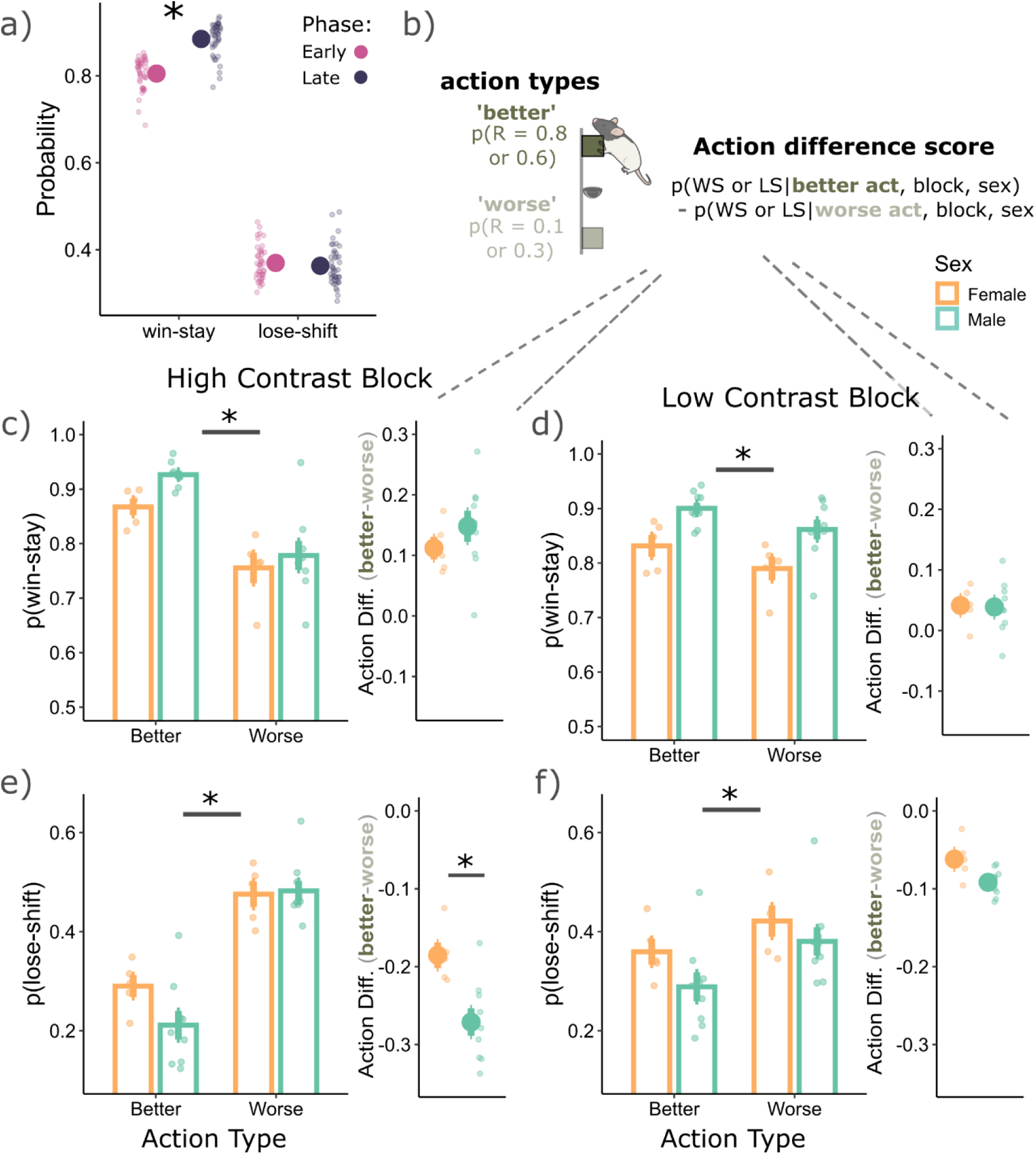
Using asymmetrical learning strategies following reversals. **a)** Rats were more likely to win-stay in the late phase of a given block (7^th^ trial until end of the block), compared to the early phase (first 6 trials of a block). **b)** To discern win and loss sensitivity between action types, we separately calculated the WS and LS probabilities for ‘better’ actions (which had a higher probability of reward), compared to ‘worse’ actions (which had a lower probability for reward). To better visualize if wins and losses were weighted asymmetrically between the two action types, we subtracted the WS and LS probabilities from the ‘better’ actions, compared to the worse. This subtraction gave an action difference score, where the further from zero, the more asymmetrically wins and losses are weighted between the two action types. In the high (**c**) and low contrast (**d**) blocks, both males and females were more likely to win-stay from better actions compared to worse (p = 0.0001, Cohen’s d = 3.23 and p = 0.01, Cohen’s d = 0.998, respectively). On the right of **c** and **d** are the action difference scores, also suggesting that better actions had a higher WS probability compared to worse. In the high (**e**) and low contrast (**f**) blocks, both males and females were less likely to lose-shift from better actions compared to worse (p = 0.0001, Cohen’s d = 8.73 and p = 0.0001, Cohen’s d = 2.93, respectively). On the right of **(e)** and **(f)** are the action difference scores, also suggesting that better actions had a lower lose-shift probability compared to worse. For specifically the high contrast blocks, males were less likely to lose-shift from better actions, compared to females (p = 0.0006, Cohen’s d = 3.30), indicating a divergence in the use of asymmetrical learning strategies between the two groups.

However, there was also a block and action type interaction for WS (F(1,36) = 16.07, p = 0.0003) and LS behaviors (F(1,36) = 107.46, p = 2.36e-12). Rats were overall more likely to use the asymmetrical learning strategy of increasing WS probability and reducing LS probability from better actions in the high contrast blocks (Figure 2c-f). This pattern was further reflected in the action difference scores (Figure 2b) adjacent to the probability figures (Figure 2c-f), where a greater difference from zero indicates more asymmetry in WS/LS probabilities between the two actions. The high contrast blocks had a larger difference in WS (F(1,13) = 22.65, p = 0.0004) and LS (F(1,13) = 175.92, p = 6.24e-09) probability between the two actions which suggest that this asymmetrical learning strategy was used more when the environment had a clearer structure.

Further, there was a main effect of sex for WS behaviors (F(1,12) = 7.23, p = 0.02). Females had a lower WS probability than males, except for the ‘worse’ actions in the high contrast block (Figure 2c). However, males and females had no significant differences in their action difference scores for WS behaviors in high (p = 0.26, Cohen’s d = 0.68; Figure 2c, right) and low contrast blocks (p = 0.93, Cohen’s d = 0.05; Figure 2d, right). This result indicates that while females did generally have a lower WS probability than males, they used the asymmetrical learning strategy of increasing the WS probability from better actions at a similar level to males. On the other hand, for LS behaviors, there was a sex and action type interaction (F(1,36) = 15.65, p = 0.0003). Males used the strategy of reducing their LS probabilities from better actions more so than females (p = 0.0006, Cohen’s d = 3.30; Figure 2e, right), specifically in the high contrast block. Therefore, specifically for LS behaviors and in the high contrast block, males and females diverged in the use of their asymmetrical learning strategies between the two action types.

Overall, males and females were more likely to WS, but less likely to LS from the better action in the late phase of both high and low contrast blocks. This result supports our interpretation that an asymmetrical learning strategy may serve the rat to remain with the ‘better’ high reward probability action, while also reducing the chances of being incorrectly influenced by the rare losses. As might be expected, we found that this asymmetrical learning strategy is used most in the high contrast block where there is a clearer task structure, and it is easier for the rat to dissociate the two actions.

### 3. Using unsigned avgRPE as a proxy for the rats’ represented uncertainty state, and the global reward state as a proxy for represented reward rate

Our behavioral results have thus far suggested that rats used asymmetrical learning strategies to adapt to the reversals in our tasks (Figures 1 and 2). Importantly, the use of these strategies was strongly modulated by the overall task structure. Rats would WS more and LS less when the optimal strategy to obtain reward was clearer, and when they had likely discerned the high reward probability action in the late phase of a block. We therefore asked what latent aspects of the task structure contributed to this observed behavior, using their choice and outcome histories. We used the unsigned avgRPE computation to represent uncertainty level and the GRS computation as a proxy for reward history (*see Methods*).

First, we aimed to establish whether the unsigned avgRPE state could reflect an aspect of the animal’s uncertainty levels. Previous uncertainty models used the current trial’s unsigned RPE, without the avgRPE component, to reflect an animal’s uncertainty state (Grossman et al., 2022; Wittmann et al., 2020). These models assume that the previous trial’s RPE does not explicitly influence the animal’s overall decision through their respective uncertainty calculations. We began by asking whether the previous trial’s RPE had any weight in the animal’s WS/LS decisions. To answer this, we fit a standard RL model, and calculated the k-value, which determines how much weight an animal placed on the current trial RPE relative to the previous trial RPE (see Equation 13) to make their WS/LS decisions. Here, a k of 1 would indicate that the rat placed complete weight on the current trial RPE, however a k of 0.8 would indicate that the rat placed a weight of 0.8 on the current trial RPE and 0.2 on the previous trial RPE (Figure 3a, also see *Methods* for details).

**Figure 3.**
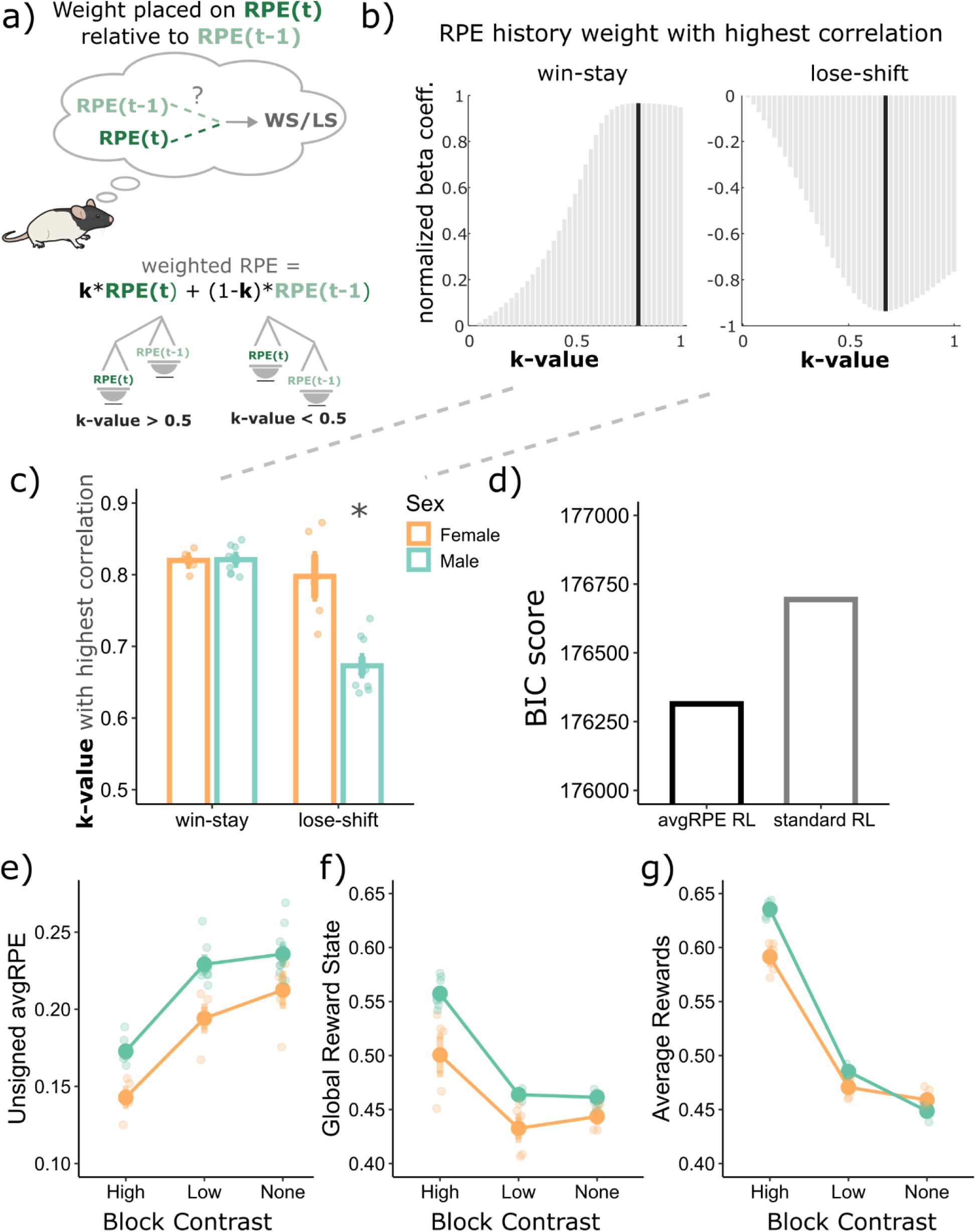
Modelling the uncertainty state as unsigned avgRPE and the reward rate as the global reward state computations. **a)** Calculating the weight placed on the current trial RPE relative to the previous trial’s RPE (k-value) when making win-stay and lose-shift decisions. A value of 1 is complete weight on only the current trial RPE, with a value of below 1 contained some weight on the previous trial’s RPE. **b)** The k-value with the highest correlation with win-stay and lose-shift behaviors was below 1, indicating that rats were influenced by the previous trial’s RPE when making their WS and LS decisions. **c)** All rats had a k-value of below 1, indicating that they were all influenced by their previous RPE when making WS/LS decisions. Additionally, males used their previous trial’s RPE more than females to make their LS decision p < 1e-04, Cohen’s d = 3.37). **d)** Bayesian Inference Criterion (BIC) score for the standard RL model which does not include an avgRPE component, compared to another model that does. The model which includes the concept of an avgRPE explained choice data better than the standard RPE model. **e)** the block with no reward contrast has the highest levels of unsigned avgRPEs, and lower for the low contrast block, with the high contrast block having the lowest levels of unsigned avgRPEs. **f)** The global reward state represents the same trend observed in empirical reward rate shown in **g.** *Abbreviations: avgRPE = average RPE, BIC = Bayesian information criterion, GRS = global reward state. * p < 0.05*.

We found that a k of close to 0.82 and 0.72 produced the strongest correlation with WS and LS behaviors, respectively (Figure 4b-c). These k-values suggest that rats use a combination of their current and previous trials’ RPE to make their WS and LS decisions in this task. There was also a main effect of sex (F(1,12) = 16.72, p = 1.5e-03) and an interaction between WS/LS factor and sex (F(1,12) = 18.6, p = 1.0e-03). We found that male rats weighted their previous trial RPE more so than females to make their LS decision (p < 1e-04, Cohen’s d = 3.37). This result is consistent with males having an overall lower LS probability, possibly suggesting that males may also need the previous trial’s RPE to be low or negative to make the decision to LS on the current trial.

**Figure 4.**
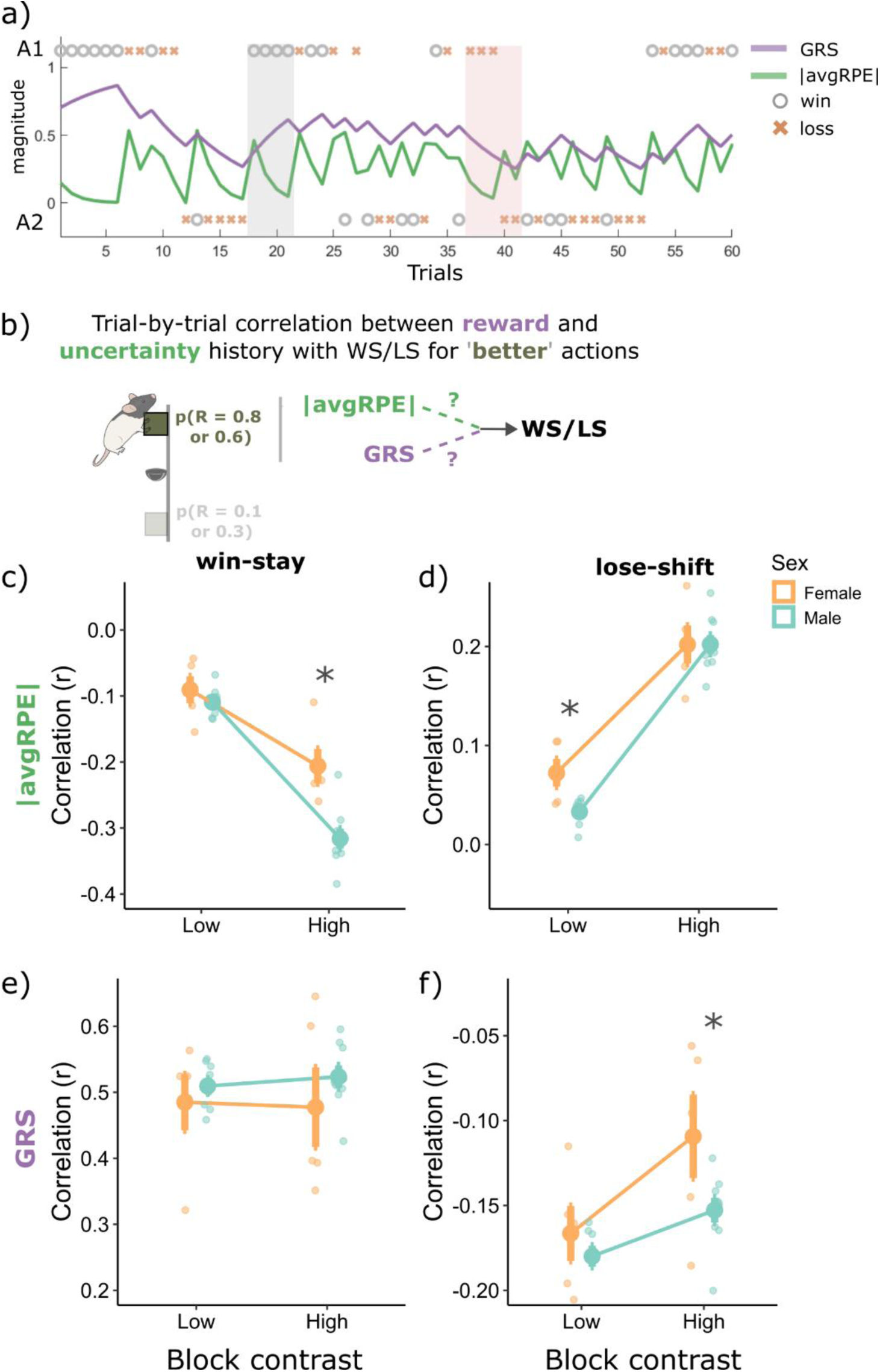
Correlations with win-stay and lose-shift behaviors from the ‘better’ high reward probability actions in the late phase, with the unsigned avgRPE and the GRS. ***a)*** an example rat’s changing trial-by-trial GRS and unsigned avgRPE trace based on the two actions and wins and losses. The grey shade highlights trials with several wins in a row, where GRS is increasing and unsigned avgRPE is decreasing. The second shade (red) highlights a series of losses, first three losses are from action 1 where GRS and unsigned avgRPE is decreasing, however, when the rat decided to shift, and take action 2 but still got a loss, the GRS continued to decrease, but unsigned avgRPE increased as this loss likely caused a negative RPE, making the rat more uncertain on the ‘correct’ action. **b)** Illustrative summary of the correlational analyses presented in **(c-d)**. **c)** Unsigned avgRPE is negatively correlated with win-stay behaviors, suggesting rats were more likely to WS from better actions when uncertainty was low. This correlation was stronger in high contrast blocks, compared to low. Interestingly, females were less influenced by the uncertainty state when making WS decision, compared to males. **d)** Rats were more likely to lose-shift from the better action when they were in a high uncertainty state, and this correlation was stronger for high contrast blocks. Interestingly, females were more influenced by their uncertainty state compared to males when making lose-shift decisions. **e)** WS behaviors positively correlated with the GRS state irrespective of the block type and sex. **f)** GRS had a negative correlation with lose-shift behaviors, and this correlation was stronger in the low contrast blocks. Therefore, both males and females, were more likely to LS from better actions when they were in a low GRS state, especially when in a low contrast block. However, the GRS influenced males’ decision more when making LS decision, as they had a higher correlation between GRS and LS decisions, compared to females. *Abbreviations: |avgRPE| = unsigned average RPE, GRS = global reward state. * p < 0.05*.

To validate whether rats are using their RPE histories, and not just their current trial’s RPE to make decisions in this task, we fit the avgRPE RL model (see *Methods*) to their behavior. We found that the avgRPE model fit the data better than a standard RL model (Figure 3c). This result suggests that a model where rats use their RPE histories captures the choice data better than a model where rats only use the current trial RPE to make their decision. Due to this better fit, we used the trial-by-trial avgRPE for subsequent analyses.

We next used this unsigned avgRPE to ask if it could be used as a proxy for the rat’s represented uncertainty state. To answer this, we examined the unsigned avgRPEs for the three different contrast blocks, with the idea that the high contrast block should have the least uncertainty (lowest levels of unsigned avgRPEs) and the block with no contrast should have the highest level of uncertainty (highest levels of unsigned avgRPEs). As can be seen in figure 3e, we found that the different block types had different levels of unsigned avgRPEs (significant main effect of block; F(2.26) = 379.6, p = 2.2e-16). Consistent with the notion that unsigned avgRPE may indicate uncertainty levels, the high contrast block had the lowest unsigned avgRPEs, followed by the low contrast block, with the block with no contrast having highest unsigned avgRPEs.

Next, we examined whether the GRS could be used as a proxy for the animal’s reward history in this task. We found consistent results between the modelled GRS (Figure 3f) and the standard average reward (Figure 3g), across the different block types. As expected, the high contrast block had the highest reward rate and GRS, suggesting that the GRS could be used as an animal’s trial-by-trial estimate of their environmental richness in this task. The GRS, like the standard reward averages, calculates reward history of a few trials back, however, unlike the reward rate, the GRS exponentially discounts previous rewards. Hence, recent rewards have a higher influence on the GRS calculation, which is not the case for the standard reward average calculation.

### 4. Uncertainty and reward history differentially influenced rats’ sensitivity to wins and losses, in a sex and block specific manner

Thus far, we found that rats used sex- and block-specific asymmetrical learning strategies in our task (Figure 1 and 2) and have demonstrated that unsigned avgRPE and GRS computations may represent aspects of uncertainty and environment richness levels, respectively (Figure 3). We next asked how avgRPE and GRS influence win and loss sensitivity, specifically for the ‘better’ high reward probability actions in the late phase, where rats were more sensitive to wins and less to losses. To answer this, we correlated trial-by-trial WS and LS, with their associated trial-by-trial unsigned avgRPEs and GRS computations (see Figure 4a for an example of the changing trial-by-trial GRS and unsigned avgRPE computations and Figure 4b for an illustration of the analysis; see *Methods* for further details).

WS behavior was inversely correlated with unsigned avgRPE for the ‘better’ action in the late phase, and this inverse relationship was stronger for the high contrast block (main effect of block; F(1,12) = 106.29, p = 2.57e-07). Rats were therefore more sensitive to wins when they were in a low uncertainty state, i.e., environments with low stochasticity. There was also a significant main effect of sex (F(1,12) = 13.87, p = 2e-03) and a sex by block interaction (F(1,12) = 8.52, p = 0.013), where males had a larger negative correlation between unsigned avgRPE and WS for the high contrast block (p = 1e-04, Cohen’s d = 2.77; Figure 4c). Therefore, the uncertainty state influenced males more so than females when making WS decisions. Next, we found that indeed rats were more likely to WS when they had a high GRS (i.e., when they were in a rich environment; Figure 4e), however, there were no main effects of block type (F(1,36) = 0.03, p = 0.87) or sex (F(1,36) = 0.90, p = 0.36). This is in sharp contrast with the unsigned avgRPE correlations with WS behaviors, showing block and sex differences. Thus, the detailed examination of trial-by-trial behavioral choices in relation to our measures of uncertainty and reward history show that estimates of uncertainty and reward density have distinctive contributions to WS behavior in this task.

LS behaviors had a positive correlation with unsigned avgRPE for these ‘better’ actions, and more so in the high contrast block, compared to low (main effect of block type; F(1,36) = 322.18, p = 2.2e-16). This positive correlation suggested that rats were more likely to maladaptively lose and shift away from the better action when their represented uncertainty state was high. Interestingly, females had a stronger negative correlation with LS and uncertainty than males (p = 0.02, Cohen’s d = 1.5; Figure 4d), which indicated that uncertainty history influenced females’ decisions to LS more so than males. Further, there was a sex by block interaction in the relationship between the GRS and LS behaviors (F(1,12) = 4.85, p = 0.048) where males were more influenced by their GRS than females when making LS decisions in the high contrast block (Figure 4f; p = 0.025, Cohen’s d = 2.52). Taken together, females were more heavily influenced by uncertainty history when making LS decisions, but in contrast, males were more influenced by the GRS when making their LS decisions. We further analyzed matching behavior between the two sex groups and found that local rewards influenced males and females’ choices similarly (see Supplementary materials Figure S2). Therefore, while overall behaviors based on rewards were generally similar between males and females, they diverged in the use of latent uncertainty and reward history computations when making these WS and LS decisions.

In sum, rats were most sensitive to wins, and least to losses from the ‘better’ high reward probability actions when their represented uncertainty was low. Further, the uncertainty state was most relevant in making WS/LS decisions when there was less stochasticity in the environment (i.e., the high contrast block), but conversely, the GRS was equally relevant for WS behaviors irrespective of environmental stochasticity, and more relevant for LS behaviors when stochasticity was high (i.e., the low contrast block; main effect of block (F1,12) = 38.20, p = 4.72e-05). These results suggest that uncertainty computations modulate behaviors more when the environment has a clearer structure, but when a clear structure is not determined, rats are more likely to simply rely on their reward histories.

Further, there were key sex differences in the influence of GRS and uncertainty history on WS/LS decisions. The uncertainty state impacted females’ decisions to lose-shift more heavily than males, however, males were more heavily influenced by their uncertainty state when making WS decisions. Conversely, both males and females were similarly influenced by the GRS when making WS decisions, however, for LS decisions, the GRS influenced males more so than females. By tracking the animal’s trial-by-trial uncertainty and reward history states, we revealed district influence of these computations on their sensitivity to wins and losses, and ultimately their behavioral strategies when adapting to changing reward contingencies in our task.

### 5. The uncertainty state and the GRS interact to asymmetrically weight wins in a block and sex specific manner

The GRS and uncertainty states differentially modulated win and loss sensitivity from the ‘better’ high reward probability actions (Figure 4) and this effect was dependent on the block type and sex group. However, it is possible that these two computations may interact, or they may be independent computations when used to modulate win/loss sensitivities. To test this, we grouped all the ‘better’ actions in the late phase (separately for the high and low contrast blocks) as high and low unsigned avgRPE state and GRS, based on a median split. We then calculated the WS and LS probabilities based on four sub-groups: 1) high unsigned avgRPE and high GRS, 2) low unsigned avgRPE and low GRS, 3) high GRS and low unsigned avgRPE, and 4) low GRS and high unsigned avgRPE. For each of these four sub-groups, we calculated the WS and LS probabilities. There were very few LS trials in the late phase of the better action, especially for the high contrast block, and as a result we focused this analysis on the WS probability’s interaction between GRS and uncertainty states only.

For males (Figure 5a), there was a main effect of the sub-group type (F(3,56) = 19.21, p = 1.09e-08), block type (F(1,36) = 8.39, p = 0.005) and a sub-group by block type interaction (F(3,56) = 3.77, p = 0.016). For the low contrast block, males would generally WS more when the GRS was high, irrespective of the uncertainty state. This result is consistent with our previous analysis in low contrast blocks where that GRS and WS have a strong correlation, but conversely, the uncertainty state and WS have a weaker correlation (Figure 4c and 4e). However, for the high contrast blocks, males did not reduce their WS behaviors when the GRS and uncertainty was low (p = 0.0001, Cohen’s d = 2.04). Instead, for these high contrast blocks, males required both a low GRS *and* a high unsigned avgRPE state to reduce their WS probability (p = 0.0001, Cohen’s d = 2.40). This result indicates that the unsigned avgRPE and GRS states may interact when modulating WS behaviors specifically in high contrast blocks in males. Conversely, females (Figure 5b) had a main effect of sub-group type (F(3,28) = 8.50, p = 0.0004), but no main effect of block (F(1,28) = 0.61, p = 0.44) or a block by sub-group type interaction (F(3,28) = 0.23, p = 0.89). Generally, a low GRS state alone, irrespective of uncertainty state, reduced the WS probability in females, and a high GRS increased this WS probability. Hence, females used the GRS more so than unsigned avgRPE to modulate their WS behaviors, which is also consistent with females having a lower correlation with unsigned avgRPE and WS behaviors, compared to males (Figure 4c).

**Figure 5.**
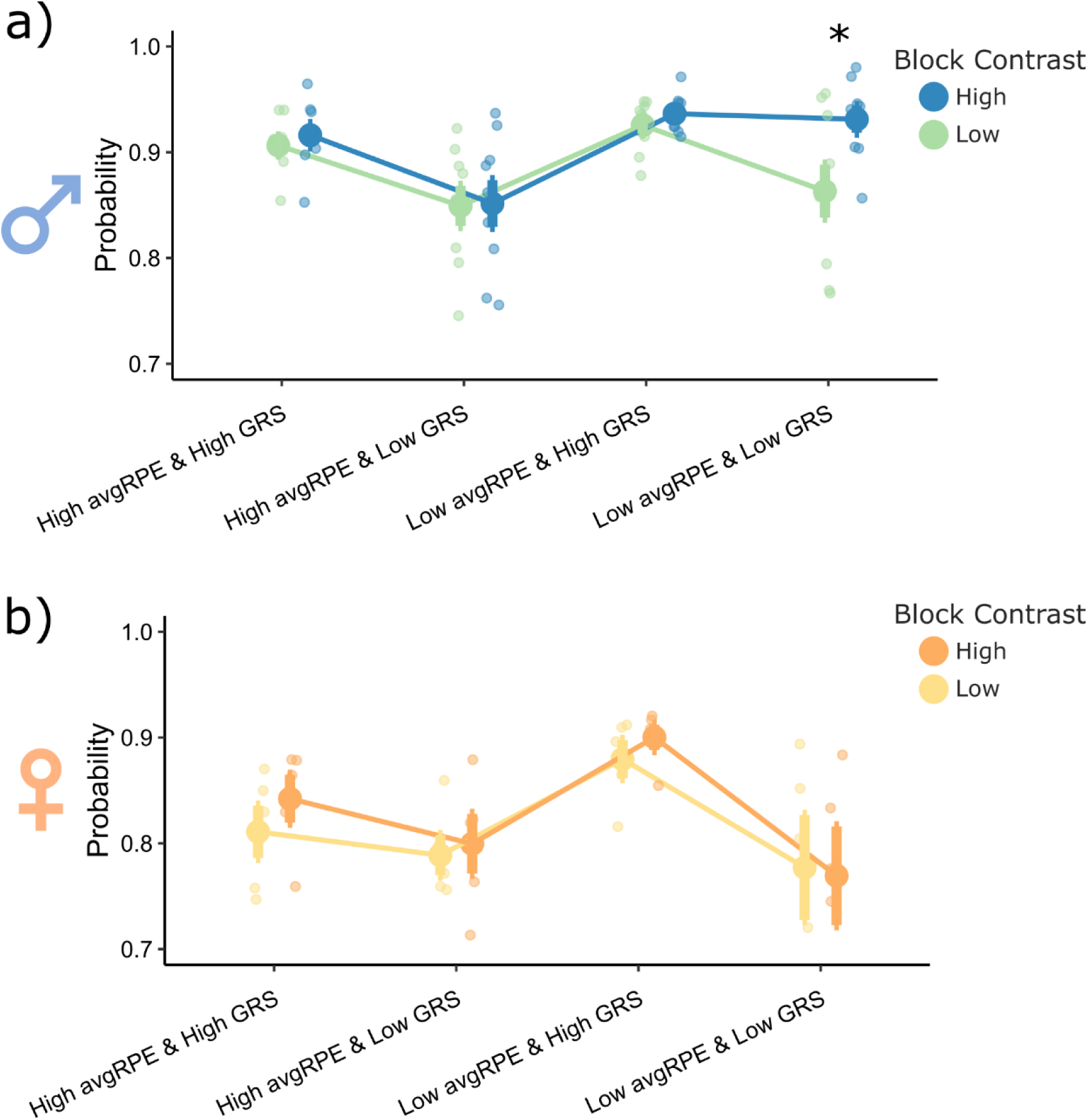
Block and sex specific interactive effects between unsigned avgRPE and GRS for win-stay behaviors from the high reward probability actions in the late phase. ***a)*** For the low contrast blocks, males would generally WS according to the GRS, irrespective of the unsigned avgRPE state. However, for the high contrast block, males showed an interaction between the two computations, where they needed both, a low GRS *and* a high unsigned avgRPE to reduce their WS probability. **b)** Females did not show this interaction effect, irrespective of the block type. Females were generally more likely to WS when the GRS was high, and less likely to WS when the GRS was low, irrespective of the unsigned avgRPE state.

Collectively, these results suggest that environmental stochasticity and sex play a role in shaping strategies involving the dynamic use of GRS and uncertainty states when modulating the weight placed on wins.

## DISCUSSION

Our primary aim was to investigate whether animals adapted to the changing levels of uncertainty and reward history by asymmetrically weighting their wins and losses. First, we established that rats indeed used asymmetrical learning strategies to adapt to the reversals in our task (Figure 1). By modelling the latent ongoing trial-by-trial uncertainty and reward history estimates (Figure 3), we found that these computations had distinct impact on rats’ WS/LS strategies (see Figure 4). Specifically, reward history, operationalized as the GRS, had an influence on rats’ decision to WS, irrespective of environmental stochasticity or sex. However, uncertainty history, operationalized as avgRPE, influenced an animal’s WS and LS behaviors particularly in environments with low stochasticity (i.e., high predictability of reward). Interestingly, uncertainty history influenced female rats’ decision to LS more than males, but conversely, male rats were more impacted by their reward history when making these LS decisions. Further, uncertainty and reward history states interacted to modulate win-sensitivity specifically in male rats and in an environment with low stochasticity (Figure 5). These results provide deeper insight into the latent computations behind the ubiquitous phenomena of asymmetrical learning and goes further to suggest that asymmetrical learning is not always a maladaptive bias but could form a part of an animal’s behavioral strategy to adapt (see Figure 6 for a schematic summary of these findings).

**Figure 6.**
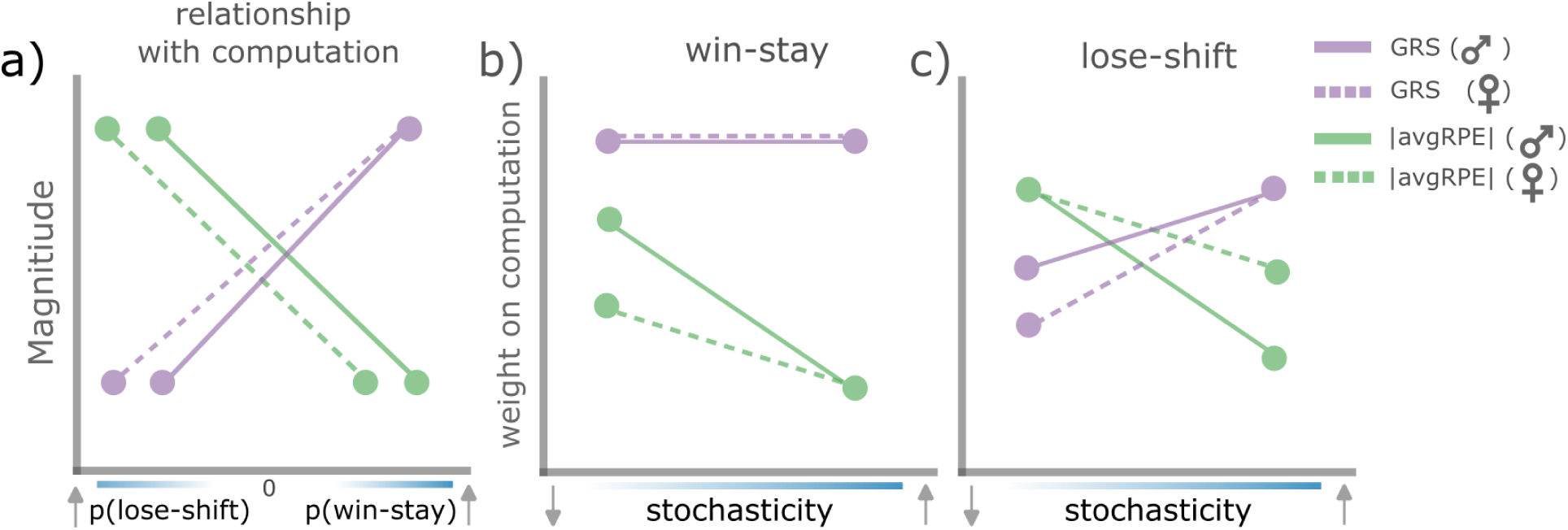
A schematic summary of uncertainty history and reward history computations when used to asymmetrically modulate win and loss sensitivities for ‘better’ actions. **a)** Both males and females LS more when the magnitude of the GRS is low, and WS more when the GRS magnitude is high. However, under the same GRS state, females are more likely to LS than males. Conversely, both sex groups overall LS more when uncertainty is high, but WS more when uncertainty is low. However, under the same uncertainty state, females are more likely to LS and less likely to WS, compared to males. **b)** both sex groups put equal weight on the GRS computation when making WS decisions, irrespective of environmental stochasticity. Under high stochasticity, both sex groups put less weight on uncertainty computations when making WS decisions, however, when stochasticity decreases (and the environment becomes more predictable), males put more weight on their uncertainty computations to make WS decisions, compared to females. **c)** Both sex groups put more weight on the GRS in an environment with high stochasticity when making WS decisions, however, in more stochastic environments, females are less influenced by the GRS than males when making LS decisions. Both sex groups put more weight on uncertainty computations, compared to the GRS when in environments with low stochasticity. However, this weight decreases in males more so than females under environments with levels of high stochasticity.

Rats adapted their behavioral strategy by increasing the win sensitivity but reducing the loss sensitivities in the later phases of the block (Figure 2), specifically for the ‘better’ action. Our interpretation here is that rats are using an exploitative strategy in the later phases, where they are more likely to have discerned which action is the high reward probability action (i.e., the ‘better’ action). Similarly, previous work has found that animals adopted an exploitative strategy in the later phases of a block or experiment (Jin et al., 2024; Mah et al., 2024; Trudel et al., 2021). And consistent with previous work (Wittmann et al., 2020), we found a positive correlation between the represented reward state (GRS) and the WS probability, and conversely, a negative correlation between GRS and LS. The positive correlation between WS and GRS can be interpreted under the optimal forging theory (Kolling et al., 2016; Wallis & Rushworth, 2014), where a rich environment (i.e., a high GRS) should make the animal more likely to stay following a rewarded choice. Our results add that this increased win-sensitivity strategy is more likely used when the environment has a clearer structure (i.e., less stochasticity, in the high contrast blocks).

However, GRS did not fully explain behavior in this task. Interestingly, we found that the represented uncertainty levels (i.e., unsigned avgRPE) differentially modulated WS and LS behaviors in our task. The represented uncertainty state had a negative correlation with WS, but a positive correlation with LS, specifically for the ‘better’ actions. Therefore, when the animal was in a low uncertainty state, they were more likely to stay following a rewarded choice from the better action. However, when that same choice was unrewarded, they were less likely to shift. The Mackintosh (1975) learning model proposed that animals gate learning in proportion to the reliability of the outcome. In the context of a low uncertainty state, a loss from this better action may be considered noise or unreliable, since this is when they are most certain they are making the ‘correct’ choice. As a result, the animal may down-weight this loss and consequently shift less. The unsigned avgRPE may be a possible computation that enables an animal to mark a loss as unreliable and thereby update less from this.

We also found key sex differences. Male rats were overall more exploitative (based on a high WS and low LS probability) than females. In contrast, previous work found that, female mice made more exploitative choices, acquiring a 2-armed restless bandit task in fewer sessions than male mice (Chen, Ebitz, et al., 2021; Chen, Knep, et al., 2021). These discrepancies may be due species differences (mice in their experiments versus rats in the current study) but could also be due to the reversal learning nature of our task, which was not the case for the previous two tasks. Another possibility is that we did not focus on task acquisition, but on strategy once the task was learnt. Future work disambiguating the sex-dependent strategies used for task acquisition versus performance may help address these discrepancies, particularly in tasks with reversals. Another paper by Aguirre et al., (2024) found that in deterministic reversal learning, but not probabilistic, males had a faster value decay of unchosen actions. Here, we add that both sex groups had similar value decays when the chosen outcome was positive, but males had a higher value decay when the chosen outcome was negative (Figure 1g-h). Collectively, these results suggest that sex differences in behaviors are significantly modulated by the task type, as well as the phase of the task (acquisition verses expert phase). We further sought to understand the possible latent computations behind differential win and loss sensitivities between the sex groups. Computational modelling suggested that the GRS was used similarly to make WS decisions between the sex groups. However, the two groups diverged in how they used their uncertainty state to modulate WS and LS decisions.

In sum, we found that indeed animals do not meet with wins and losses the same. However, this asymmetry was a part of their behavioral strategy to adapt. We found that uncertainty and reward history may underlie this common phenomenon of asymmetrical learning, allowing the animal to adapt appropriately to the changing environment. While we show that asymmetrical learning can be used adaptively, there are cases where this process can go awry and produce decision-making pathologies, including addiction (Kalhan et al., 2021, 2022, 2024), depression (Ubl et al., 2014), post-traumatic stress disorder (Morey et al., 2008, 2015) and obsessive compulsive disorder (Fitzgerald et al., 2005; Suzuki et al., 2023). Our results have implications for better understanding these decision-making pathologies and suggest that a misalignment of uncertainty and reward history computations may contribute to some aspects of maladaptive asymmetrical learning symptoms. Further, while it is known that reward history and uncertainty computations produce adaptive behaviors, we add that asymmetrically modulating wins and losses may be one possible mechanism by which this is achieved. Hence, future models of learning and decision-making that can incorporate this additional computation may help better explain decision-making processes.

## METHODS

### 1. Subjects

We used a total of 28 Long-Evans rats (n = 18 males, n = 10 females; 10 weeks old upon arrival, Envigo). They were all single housed and kept in a constant temperature and humidity-controlled environment, with a 12-hour light/dark cycle (light cycle starting 7am, and dark cycle starting at 7pm). All experiments were conducted during their light cycle.

### 2. Ethics Statement

The experimental procedures were done in accordance to the Johns Hopkins University Animal Care and Use committee. The protocol number is RA23A232.

### 3. Behavioral task

#### 3.1. Initial training

Behavioral training and testing were done in a customized operant chamber (Med Associates). During both, training and testing periods, rats were water restricted while maintaining at least 90% of their *ad-libitum* weight. The first step of training involved learning to enter the reward magazine to collect a reward (100μL of 10% sucrose with tap water). Rewards were delivered randomly in the magazine, but with an average interval of 60 seconds. Next, rats learnt to initiate a trial through this magazine entry, which then triggered the insertion of two levers at the same wall as the magazine, with one lever to the left, and the second to the right of the magazine. At this stage, pressing any lever delivered a reward in the magazine, with 100% probability. Following a successful completion of lever press training, rats performed a simple reversal learning task. Here, pressing one of the two levers led to a reward with 100% probability, and the other lever a 0% probability of reward. The reward probabilities for the levers were reversed (the previously rewarded lever press now became the unrewarded press, and vice versa for the previously unrewarded lever press) randomly after passing a performance threshold of 0.75 (calculated as an exponential moving average of the previous 8 trials). Hence, the reward probabilities reversed following a successful acquisition of the correct lever press to obtain a reward. Here, the reversal probability after reaching the threshold was set at 10%, and if a reversal did not happen within 20 trials of surpassing the performance threshold, a reversal was forced. This deterministic reversal learning training lasted between 1 to 3 days. Following this, the probabilistic reversal learning task was started (see below).

#### 3.2. Probabilistic reversal learning (PRL) task

The PRL task consisted of one of the two levers having a 70% probability of being rewarded given a press of this lever, and the second lever being 10% reward probability given a press of this lever. The reversal rules were identical to the deterministic reversal learning task described above. The trial was signaled as ready to be initiated by a magazine light on, and when the rat entered the magazine, the two levers inserted after a variable delay (100ms-1000ms, with randomly selected intervals of 100m). If rats did not press a lever within 30s, this was termed as an invalid trial, and this trial was excluded from further analysis (less than 1% of trials for all rats). Immediately upon lever press, rewarded trials were indicated by two clicker sounds (0.1s interval, generated by MED Associates, ENV-135M), with unrewarded trials cued by 0.5s of white noise (generated by MED Associates, ENV-225SM). Reward consisted of ∼33μL of the sucrose solution, delivered in the magazine via a syringe pump (MED Associates, PHM-100). A rewarded trial was termed complete after the first exit by the rat from the magazine, having collected the reward. An unrewarded trial ended after the white noise cue presentation. The intertrial interval was 2.5s. All rats received a total of 15 2-hr sessions. See Figure 1 for the basic task structure, and Figure S1 (supplementary materials) for the data from this task.

#### 3.3. Dynamic probabilistic reversal learning (DynaPRL) task

Fourteen rats (9 males, 5 females) from the PRL task advanced to training on the dynaPRL task. The overall task structure of the dynaPRL task is identical to the PRL task, with three important differences. First, the reward probabilities of the two actions were different. The dynaPRL task consisted of three block types (distinguished by their reward probability, given the lever press). First, one block type had a 45% reward probability if either of the two levers were pressed (termed a no action contrast (NC) block). The second block type was a 60% reward probability for one lever press, and 10% for the other (termed low action contrast (LC) block). The third block type was 80% reward probability after pressing one lever, and 10% for pressing the other (termed high action contrast (HC) block). Importantly, each block diverged in the difference – i.e., the contrast between the two reward probabilities, hence labeled as the high, low and no contrast blocks (Figure 1b). The second difference between the PRL and dynaPRL tasks was when the reward probability contingences switched (i.e., block transitions). Here, a block switched every 15-30 trials, with the exception of when rats made at least four consecutive incorrect choices in the blocks with an action contrast (LC and HC blocks). The block transitions occurred randomly, but with two rules imposed, 1) neutral blocks were not repeated, and 2) the position of the lever with the higher reward probability was not repeated. The third key difference between the two tasks was that the dynaPRL task also had a reward magnitude manipulation, but only in the HC and LC block types. Specifically, after 12 trials within a block had passed, in 40% of these trials, the reward was either halved (16μL) or doubled (66μL). Rats performed an average of 30 sessions, with each session lasting 2 hours.

### 4. Analyses

#### 4.1. Model-independent analyses

All the model-independent analyses were focused on calculating the win-stay (WS) and lose-shift (LS) probabilities. An action was termed WS if that action was rewarded, and then was subsequently repeated. An action was termed LS if an action was not rewarded, and the subsequent action was changed. In general, WS probabilities were calculated as the number of WSs, divided by the number of wins in the trials of interest. Similarly, the LS probability was calculated as the number of LSs, divided by the number of losses, in the trials of interests.

In subsequent analyses we calculated the WS and LS probabilities separately for early and late phases of a given block (Figure 2). The early phase included the first 6 trials after a block transition, and the late phase extended from the 7^th^ trial until the end of the block. Therefore, here, we calculated the total number of WSs or LSs in a given phase of a given block, and divided by the number of wins or loses within that block/phase. For example, when calculating the WS probabilities of the early phase of the HC block, we first calculated the number of WSs in the first six trials of this block for every session. We then summed all the number of WSs from every session in the early phase of this HC block. Next, we calculated the number of wins in the early phase of every high contrast block, in every session. This was then summed. Therefore, to calculate the probability of WS in the early phase of the high contrast block, we divided the number of WSs here, divided by the total number of wins here. The same analysis was used for the other block types, and for the late phase.

We also did an analysis where WS and LS probabilities were calculated separately given the action type (high or low reward probability action), the block type (HC, LC and no contrast) (Figure 2). Here, a similar principle was applied as on the above analyses, but this time also taking into account the action type. For example, the probability of WS from the high reward probability action in the early phase of the high contrast block was calculated as following. For each session, we first calculated the number if WSs given the high reward probability action was made, and in the early phase of the HC block. We then summed this across all sessions, to get the total number of WSs given the high reward probability action was made in this block and phase type. Next, we calculated the number of wins from this high reward probability action in this phase and block type, of a given session, and then summed this across all sessions. The WS probability was therefore calculated as the number of WSs from the high reward probability action in the early phase of the HC block, divided by the number of wins from this action/phase and block type. The probability of WS and LS was calculated similarly for all the block, phase and action types.

Lastly, we calculated the p(preservation) (i.e., the probability of persisting the with same choice) pre- and post-reversal to asses adaptation to the reversals in our task (Figure 1c). This p(preservation) score was calculated trial-by-trial using the ratio of correct actions (i.e., probability of the correct choice) and the total number of blocks. However, in the no contrast block, there is no ‘correct’ choice, and we therefore randomly selected one of the actions as ‘correct’ for the calculation. There were five trials in the pre-reversal stage, and twelve trials post-reversal.

#### 4.2. Model dependent analyses

##### 4.2.1. Models used

We used four different models in our analyses. The first model was a standard reinforcement learning (RL) model which consisted of the learning rate (α) and the inverse temperature (β) as free parameters. Here, the trial-by-trial reward prediction error (RPE) was calculated as:

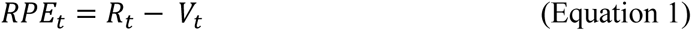

Where t is the trial, R is the reward and V is the value. The value was then updated as the following:

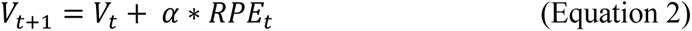

We used a softmax function mechanism to calculate the choice probability of the two actions, dependent on the value of the two competing actions:

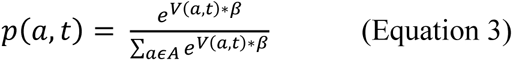

Where *p*(*a*, *t*) is the probability of taking action a, at trial t, and β is the inverse temperature (where low values give close to random action selection, and higher values increases the probability of selecting the action with the highest values).

The second model used was the average RPE (avgRPE) RL model. Here, we calculated the RPE as in equation 1 above, and using the RPE from here, we calculated the exponential-like avgRPE as below:

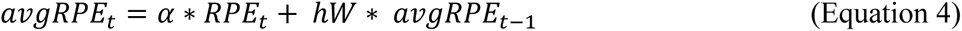

Where *α* is the learning rate and, hW is the history weight which was a free parameter, bounded between 0 and 0.75. A high hW indicated more weight placed on the previous avgRPE, and conversely, a low value here indicated a lower weight placed on the previous avgRPE. The value was then updated as below:

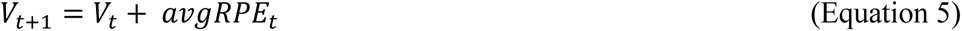

Importantly, if hW is equal to 1, the model becomes the classic RL model (model 1). The choice probability calculation used was identical to the classic RL model as above, using the softmax action selection equation (equation 3). This avgRPE RL model therefore had three free parameters (α, β and hW).

The third model used was the global reward state model (GRS), also used by (Wittmann et al., 2020). Here, the reward trace (R-trace) was calculated as an exponential moving average of the rewards as below:

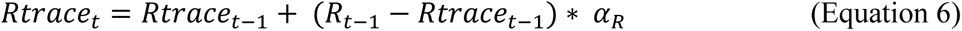

Here, *α*_*R*_ is the weight placed on the reward history, and was bounded between 0 and 1. Using this R-trace, the RPE was then calculated as below:

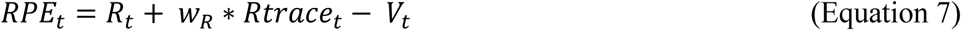

Here, *w*_*R*_ was the weight placed on the R-trace calculation and was bounded between -1 and 1. The softmax action selection mechanism was used (equation 3), and the value was updated as in equation 2 above, however, this time using the RPE from equation 7. Overall, this third GRS model had 4 free parameters (α, β, *w*_*R*_ and *α*_*R*_).

The fourth model was a five parameter RL model, used specifically to understand general asymmetrical learning differences between the two sex groups. This model had two learning rate parameters (one for learning from positive outcomes; α+, and a second for learning from negative outcomes; α-). The next two free parameters were value decay rates from the unchosen actions (one decay rate for the unchosen positive outcome; γ+, and second for unchosen negative outcome; γ-). The 5^th^ free parameter was the inverse temperature (β), used to select an action based on softmax action selection (Equation 3). The value in this model was updated as following:

At trial t, if the chosen action is A1, and this chosen action was rewarded (R = 1), the value of the chosen action (A1), and unchosen action (A2) was updated as:

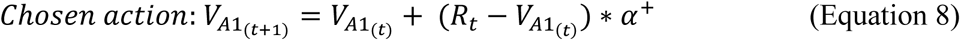

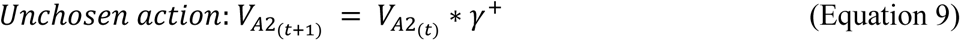

If the chosen action (A1) was unrewarded (R=0) at trial t, the value was instead updated as:

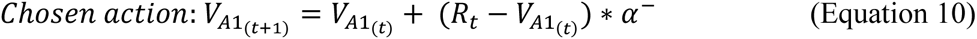

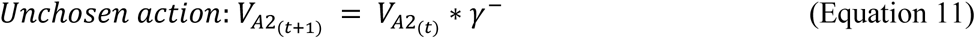

##### 4.2.2. Model fitting and comparisons

All models, except the five parameter RL model, were fit on a session level, using the MTLAB (2024b) function *fmincon*, to find parameters that maximized the log-likelihood. Each session fit was repeated 10 times, and the parameters that yielded the highest log-likelihoods were then used for further analyses. Model comparison was done based on the Bayesian Information Criterion (BIC) score, calculated on a session level as:

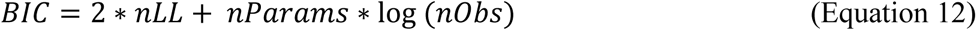

Where, nLL is the negative log likelihood, nParams is the number of free parameters, and nObs is the number of observations.

Given that the five-parameter RL model had a relatively large number of free parameters and it specifically was used to compare the parameter estimates between the two sex groups, we used hierarchical model fitting and comparisons at the group-level such that the likelihood of over-fitting due to noise between individual rats is reduced (Carpenter et al., 2017). Here, we constructed group-level (male and female) hyperparameters that then modulated the estimates for the five parameters for each rat in a given group. This procedure was done using the Matlab Stan interface (https://mc-stan.org/users/interfaces/matlab-stan). We estimated the best fitted model parameters using the overall choice and outcome data for each rat, per session. More specifically, we drew consecutive samples from the posterior density of the model parameters using Hamiltonian Markov chain Monte Carlo – which gave the density estimates for the best fitted model parameters per rat. Importantly, each rat had the parameter estimates for their five free parameters, all of which were drawn from the group-level hyperparameters (group mean (μ), group mean difference (δ), and variance (σ). In order to asses the magnitude of the differences between the parameter estimates between groups, we used the directed Bayes Factor (dBF) – which calculated the ratio of proportion of the group difference hyperparameter (δ) above zero, versus the proportion that is below zero. The larger the dBF magnitude, the greater the group differences between the parameter estimates of interest. We have previously used this model fitting approach, see Cheng et al., (2025) for more details on this method.

#### 4.3. Model-dependent correlation analyses with WS-LS behaviors

To discern whether rats only use the current trial RPE or a combination of the current and previous trial’s RPE in order to make their WS/LS decisions (Figure 5b), we used the equation below:

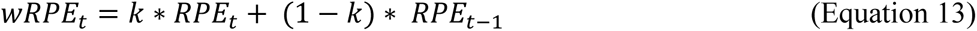

Where, wRPE is the weighted RPE based on the current and previous trial RPE. The RPEs here were estimated based on the classic RL model (see above). The k-value determined how much weight to place on the current trial RPE relative to the previous trial. A k of 1 would only place the weight on the current trial RPE, however a k of 0.8 would place a weight of 0.8 on the current trial RPE and 0.2 on the previous trial RPE. We therefore used equation 9, with k of between 1 and 0.5, in increments of 0.025, to generate trial-by-trial, wRPEs. After generating wRPEs with different ks, we used binomial logistic regression to correlate all the wRPEs (that were z-scored) with the rats’ WS and LS behaviors to determine the k-value that produced the strongest correlation (i.e., beta weight).

#### 4.4. Statistics

All statistical analyses were done using a linear mixed effect model, using RStudio 4.1.2 (function “*lmer*”). In each case we input the factors of interest for that analysis (main effects and interactions), as specified in the results section for each analysis. Further, to account for the variability due to individual rats, we used the rat ID as a random effect in all our liner mixed effect models. The effect size was calculated as the Partial eta-squared (η_p_^2^), with 0.01 indicating a small effect size, 0.06 indicating a medium effect size, and 0.14 indicating a large effect size. Where the linear mixed effect models yielded significant results, post-hoc analyses were done using t-test comparisons (function *“emmeans”*), adjusted for multiple comparisons using the Tukey method. To determine the effect size of the t-tests, we used Cohen’s d, where 0.2 indicated a small effect size, 0.5 indicated a medium effect size, and 0.8 indicated a large effect size.

## Acknowledgments

Supported by NIH R01AA031609 (PHJ), a Brain & Behavior Research Foundation Young Investigator Grant (YC), and a JHU Kavli NDI Fellowship (YC). We would also like to thank Professor Tom Verguts for helpful discussions.

## Author Contributions

SK: conceptualization, data curation, formal analysis, methodology, software, visualization, writing – original draft preparation, writing – review and editing. RM: investigation. YC: investigation, methodology, conceptualization, writing – review and editing. PHJ: conceptualization, methodology, funding acquisition, supervision, writing – review and editing.

## Data availability statement

Data will be made available in the Open Science Framework repository upon publication.

## Supplementary Materials

**Figure S1.**
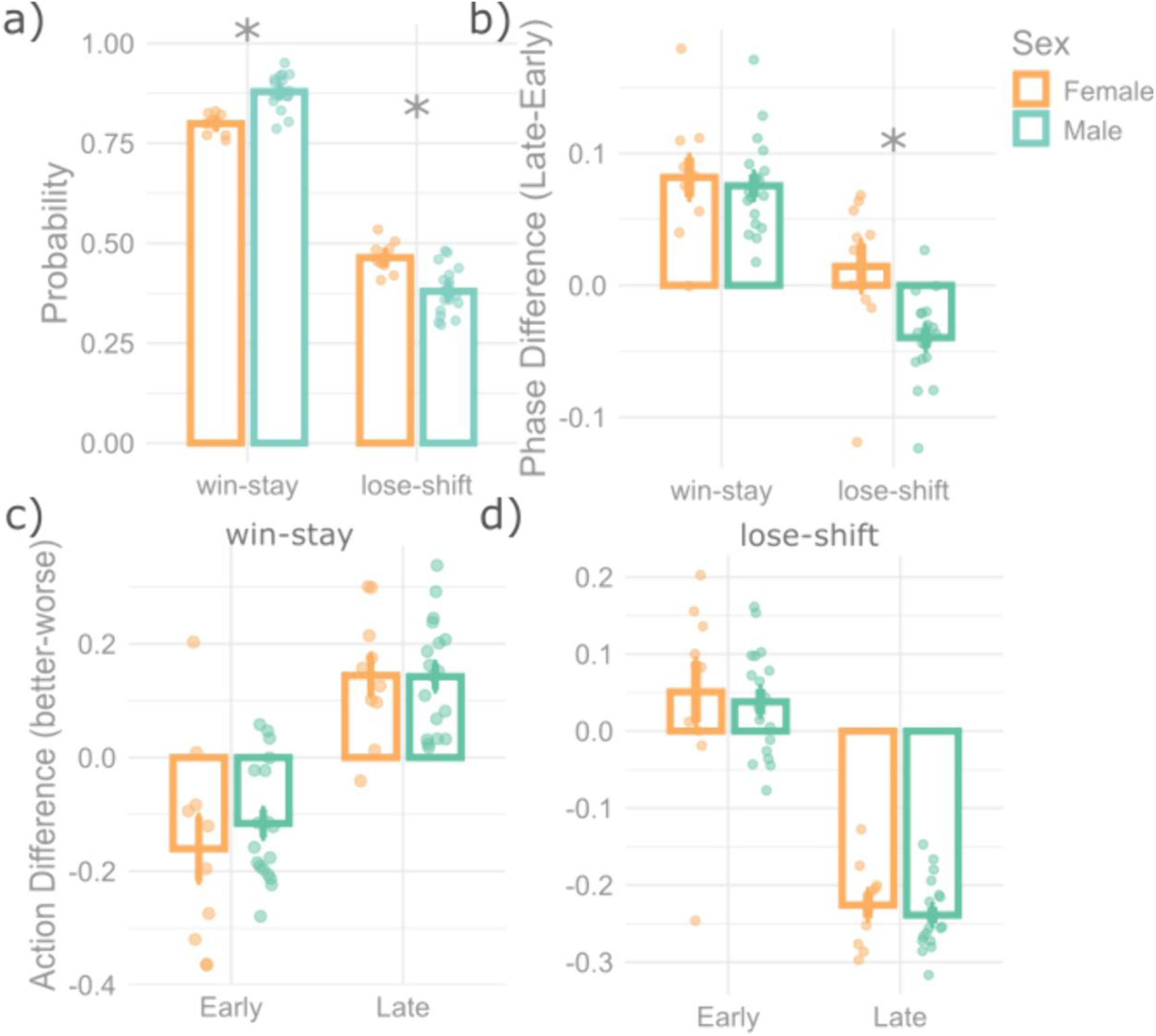
Asymmetrical learning strategies in the probabilistic reversal learning task. a) rats had a higher WS probability compared to the LS probability in the PRL task (main effect; F(1,52)=1076.6; p = 2.2e-16). There was an interaction between the WSLS factor and sex (F(1,52) = 41.83, p = 3.5e-08). Subsequent t-tests revealed that, females had a lower WS probability, but a higher LS probability, compared to males. **b)** asymmetry emerges for WS and LS behaviors (main effect of WSLS factor; F(1,26) = 66.55, p = 1.2e-08), where both males and females WS more in the late phase of the block, compared to the early phase. However, only males developed the strategy of also reducing their LS probability in the late phase of the block (p = 0.002, Cohen’s d = 1.34). **c-d)** There were differences in WS and LS probabilities between phases (interaction between WSLS type and phase type; F(1,104) = 198.86, p = 2.2e-16). Subsequent t-tests revealed a significant difference in early versus late phase WS (p < 1e-04, Cohen’s d = 2.80) and LS probabilities (p < 1e-04, Cohen’s d = 2.76). In the late phase, rats were more likely to WS if that win was from the better action, compared to the worse action. Conversely, rats were less likely to LS if that loss was from this better action, compared to the worse action, which emerged in the late phase of the block.

**Figure S2.**
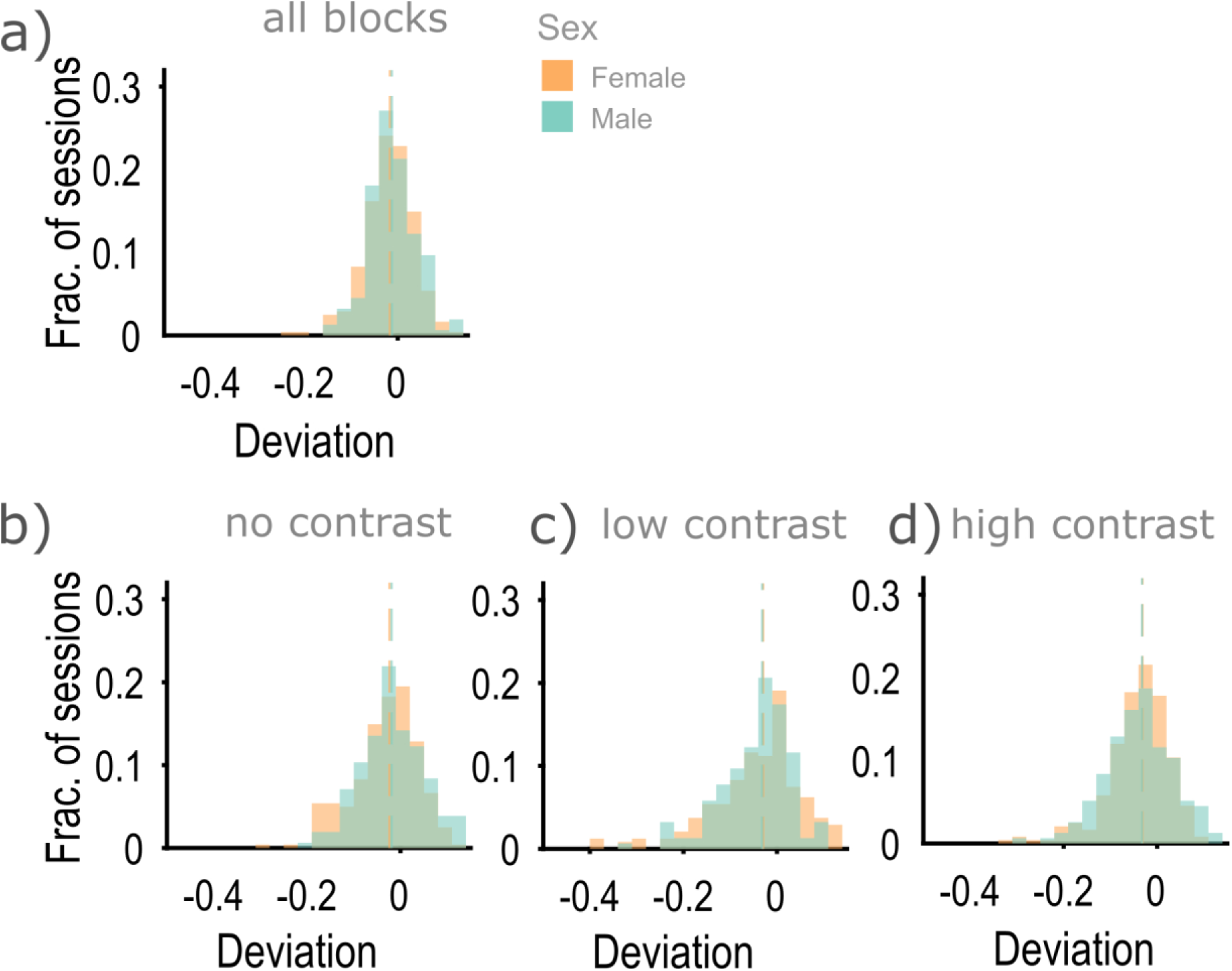
Males and females deviate at similar levels from the matching behaviors. **a)** Both sex groups have a higher matching score (i.e., a higher deviation of choice probability from the reward probability based on the choice). However, in across all blocks **(a),** and within the three block types **(b-d)**, there were no sex differences, indicating that males and females make similar choices based on their local reward outcomes.

